# Paradoxical relationships between active transport and global protein distributions in neurons

**DOI:** 10.1101/2020.08.05.238071

**Authors:** A. Bellotti, J. Murphy, L. Lin, R. Petralia, Y-X Wang, D. Hoffman, T. O’Leary

## Abstract

Neural function depends on continual synthesis and targeted trafficking of intracellular components, including ion channel proteins. The detailed biophysics active ion channel transport are increasingly well understood, along with the steady-state distribution of functional channels in the membrane. However we lack a quantitative understanding of how transport mechanisms give rise to stable expression patterns, and how live measurements of active transport relate to static estimates of channel density in neurites. We experimentally measured neuronal transport and expression densities of Kv4.2, a voltage-gated transient potassium channel. Kv4.2 is known to have a highly specific dendritic expression and little or no reported functional expression in axons. Surprisingly, in over 500 hours of quantitative live imaging, we found substantially higher microtubule-based transport of Kv4.2 subunits in axons compared to dendrites. We show that this paradoxical result is expected using a mass action trafficking model of intracellular transport that we calibrate to experimental measurements. Furthermore, we find qualitative differences in axonal and dendritic active transport that are captured in a stochastic model of puncta transport. This reveals that active transport is tuned to efficiently move cargo through axons while promoting mixing in dendrites. Finally, our data reveals trends in transport parameters that can explain the functional density profile of Kv4.2. Puncta velocity bias is directed distally and the magnitude of this bias increases with distance from the soma. These trends are consistent with an analytical solution of a linear transport PDE, corroborating previously unexplained distributions of Kv4.2 subunit localization and A-type current density. Together, our results provide new quantitative data on ion channel trafficking and reveal counterintuitive but mathematically consistent relationships between the distribution of cargo that is in transit and its functional expression.

**SIGNIFICANCE:** This study of ion channel transport reveals a seemingly counterintuitive result: the majority of subunit transport occurs in axons for a cargo whose static distribution is concentrated in dendrites. This disparity is reconciled by a simple mathematical model of transport, which reveals that the local density of actively transported intracellular cargo can show an inverse relationship with its static expression density. Mass action models also reconcile the previously unexplained, highly asymmetric, increasing distribution of Kv4.2 with its measured trafficking density that resembles diffusion with minimal drift. The generality of our analysis prompts caution in how static snapshots of intracellular cargo distributions should be interpreted for any type of intracellular cargo.

## INTRODUCTION

Most cell types exhibit polarity, an intrinsic asymmetry that defines subcellular directionality and compartment identity. Neurons exhibit strong polarity, with most neuron types showing clear morphological and functional differences between axons and dendrites that shapes the flow of information. The functional differences between these compartments is underpinned by localization of proteins, organelles, and other material within these cell projections. Such global patterning of intracellular expression is thought to be largely mediated by intracellular transport (1, 2) which is increasingly well understood. However, we still lack an understanding of how intracellular transport gives rise to consistent expression profiles in complex neuronal morphologies (3–5).

Kv4.2 is a voltage-gated ion channel prevalently expressed in excitatory neuron dendrites. Kv4.2 conducts A-type, transient outward potassium current, which is abundant in dendrites but scarce in axons (6). Dendritic expression of Kv4.2 is consistent with its hypothesized role in dendritic integration and control of excitability (4, 6, 7). Moreover, A-type current exhibits a five- to sixfold increase in current density over the length of the apical dendrite (7). Localization studies of Kv4.2 corroborate this finding, showing a 70% increase in channel density along the apical dendrite (8). Kv4.2-mediated A-currents have not been reported in axons, but other channels that pass A-current have been found in axons (6, 9, 10). Effective axonal surface expression of Kv4.2 therefore seems unlikely.

However, the reported amount of Kv4.2 subunits localized in axons varies substantially among quantitative localization studies. Alfaro-Ruíz et al report only 1.2% of total CA1 immunogold particles are found in axons (11). Kerti et al contrastingly report nearly 20%, and the authors remark that “[this result] is surprising, because the Kv4.2 subunit is conceived as a somato-dendritic ion channel” (8). In this study, we found predominant endogenous expression of Kv4.2 in dendrites with a non-negligible presence in axons. Due to the imaging modality, these studies cannot establish whether subunits present in axons are in transit or whether they are in membrane-bound, functional channels.

Several studies have measured Kv4.2 trafficking and internalization in dendrites (12–15), but none to date have enabled a quantitative, global model of expression patterns or sought to account for the presence in axons of a subunit whose functional expression seems confined to dendrites. Surprisingly, during live imaging of transfected, actively transported subunits, we found a greater density of microtubule-based transport of Kv4.2 in axons than in dendrites. This paradox of unexpectedly high subunit density in regions of low functional expression turns out to be consistent with a simple mass action model of intracellular transport. Assuming higher propensity for surface transport in dendrites, the model predicts a resultant lower density of microtubule-bound cargo; conversely, a low probability of surface transport in axons results in more cargo remaining on the microtubule.

Our model accommodates and accounts for qualitative differences in microtubule transport between axons and dendrites, including displacement, directional bias, speed, and stall time. We inferred these parameters for axonal and dendritic puncta and found that axonal transport exhibits higher state dependence - or memory - enabling longer uninterrupted runs and faster transport. Dendritic transport parameters, by contrast, show higher diffusivity, consistent with a mechanism that provides opportunity for membrane trafficking mechanisms to interact with trafficked subunits.

Finally, we used our model and experimental measurements to account for an increasing functional Kv4.2 density from proximal to distal compartments. This expression pattern has been extensively characterized and is important for dendritic function, but the question of how it emerges from a relatively simple trafficking mechanism has remained unanswered. We experimentally estimated model parameters including microtubule occupancy and transport rates as functions of distance from soma. Constrained with these data, we provide an analytical solution for the microtubule occupancy profile that can recapitulate the previously unexplained Kv4.2 localization profile along dendrites.

## MATERIALS AND METHODS

### Animals and cell culture

All animal procedures are conducted with accordance of the National Institutes of Health Guide for the Care and Use of Laboratory Animals under a protocol approved by the National Institutes of Child Health and Human Development’s Animal Care and Use Committee.

#### Rat hippocampal dispersed cultures

Hippocampal cultures are prepared from gestational day 18-19 wildtype Sprague-Dawley rats as previously described (14). Briefly, fetal pups are removed from the mother and hippocampus tissues are dissected and placed in dissection media (DM). For 500 ml of DM, we filter sterilized: 50 ml 10x HBSS (Gibco™, 14185-052), 5 ml pen/strep (Gibco™, 15140122), 5 ml pyruvate (Gibco™, 11360070), 5 ml HEPES (1M, Gibco™, 15630080), 15 ml of 1 M stock solution glucose (from powder, Sigma), 420 ml Ultra Pure Water (Kd Medical Inc).

Tissue was mixed with papain (Worthington Biochemical Corp., Lakewood, NJ) for 45 min at room temperature. Tissues were rinsed for removal of extracellular material with dissection media several times, and dissociated cells were plated in Neurobasal™ media (ThermoFisher, Waltham, MA) with 5% FBS (HyClone Characterized Fetal Bovine Serum, SH30071.03, GE Healthcare LifeSciences, Pittsburgh, PA), 2% GlutaMAX (ThermoFisher), and and 2% Gibco™ B-27™ supplement (ThermoFisher) (subsequently called NB5 media). Cells were incubated in 5% CO_2_ at 37 deg C. After 24 hrs, cells were transferred to Neurobasal™ containing 2% GlutaMAX, 2% Gibco™ B-27™ supplement (NB0 media). Half of the media is replaced with fresh NB0 media every three to four days, and cells are imaged after 9-13 days *in vitro*.

#### Construct

A Kv4.2 construct was conjugated at the N-terminus to strongly enhanced green fluorescent protein (SGFP2) (16), henceforth referred to as Kv4.2-SGFP2. pSGFP2-C1 was a gift from Dorus Gadella (Addgene plasmid # 22881; http://n2t.net/addgene:22881; RRID:Addgene_22881). We sub-cloned mouse Kv4.2 into the SGFP2 plasmid using NheI amd SalI restriction sites.

#### Transfection

Lipofectamine^®^ 2000 transfection was performed following manufacturer protocol with some modifications. 2 μl of Lipofec-tamine^®^ 2000 Transfection Reagent (ThermoFisher) and 2 μg of DNA plasmid were each diluted in 200 μl of Neurobasal™ media and incubated at room temperature for 5 min. The two solutions were then combined and incubated at room temperature for 15-20 min. 100 μl of total mixture was added to each well and incubated at 37 deg C for 4 hrs before changing media. The cells were then incubated for an additional minimum of 1 hour before imaging.

#### Immunostaining

Following hour-long time series, samples reserved for antibody staining were fixed/permeabilized and immunostained as previously described (17, 18) and briefly reiterated here. Upon completion of time series, the coverslips were removed from imaging chamber and the location of the neuron of interest was labeled with a fine tip marker. Coverslips were fixed with 4% paraformaldehyde (Electron Microscopy Sciences, R 15710, Hatfield, PA) with 4% sucrose (Sigma, S9378) at room temperature for 15 min followed by three 1X PBS (Gibco™, 14190) washes before overnight storage in 1X PBS at 4 deg C. Coverslips were permeabilized in 0.2% Triton X-100 (Sigma, T8787) for five minutes at room temperature and washed once in 1X PBS for 5 min. Cells were incubated for one hour at room temperature in 0.04% Triton X-100 solution in 1X PBS containing 1:100 dilution of anti-ankyrin-G rabbit primary antibody (NeuroMab, 75-146, Davis, CA) or 1:1000 dilution of MAP2 antibody (Chemicon, Burlington, MA). Upon completion of primary incubation, coverslips are washed three times with 1X PBS for 5 min. Coverslips are then incubated with secondary antibodies anti-rabbit-555 (1:500) for ankyrin-G or MAP2 and anti-GFP-488 (1:400) (Molecular Probes, Eugene, OR) for one hour at room temperature before another three washes with 1X PBS. Coverslips were then mounted onto glass slides using ProLong™ Diamond Antifade Mountant containing DAPI (Invitrogen, Carlsbad, CA).

### Microscopy

#### Hour-long FRAP recordings

18-mm coverslips were removed from wells and placed in a Quick Release Chamber (Warner Instruments, QR-41LP, 64-1944, Hamden, CT). Cells were immersed in 800 μl imaging buffer consisting of 1x Tyrode’s solution: 135 mM NaCl, 5 mM KCl, 2 mM CaCl_2_, 1 mM MgCl_2_, 25 mM HEPES, 10 mM glucose (all Sigma-Aldrich) at pH 7.4. All imaging was carried out at the NICHD Microscopy and Imaging Core using a Zeiss LSM 710 laser scanning confocal microscope (Carl Zeiss Microscopy LLC, White Plains, NY). Global still images were captured using a 40X oil-immersion objective and stitched together in ImageJ.

Fluorescence recovery after photobleaching (FRAP) time series was performed using a 63X oil-immersion objective with the pinhole diameter set to 4 airy units. The 495-nm laser line was used for both imaging and bleaching. During imaging, laser power was set to 4%, and during bleaching power was set to 100%. Images were acquired at 1024 × 1024 resolution at 1.0X optical zoom with 750x gain. Time-lapse images were captured at 0.2 Hz for 60 to 85 min using Zeiss LSM Image Browser software. Z-plane focus was maintained using Zeiss Definite Focus after each frame captured. The cells were temperature- and CO_2_-controlled at 37 deg C and 5% during imaging using a stage top incubator (Tokai Hit, STXG-WSKMX-SET, Fujinomiya, Japan). Every 10-20 minutes, Kv4.2-SGFP2 was bleached to 30-70% of baseline intensity by the same 495-nm laser (100% power, 10 iterations) over the bleach region of interest (ROI).

#### Extended FRAP recordings

Coverslips were maintained for extended recordings of up to 11 hrs. A distilled water reservoir was placed adjacent to the imaging chamber, and the chamber was covered with a 35-mm Petri dish to maintain humidity. Time-lapse images were captured at 0.1 Hz with recurrent photobleaching every 50 min. All other procedures and conditions are as previously described.

#### Colchicine treatment

Samples treated with colchicine (Sigma, C9754) were prepared and time series were captured as previously described for hour-long recordings. At 30 minutes into the recording, 1 μl DMSO (control) or 1 μl DMSO containing 40 μg colchicine was mixed into the imaging chamber, for a final colchicine concentration of 125 μM. Number of mobile puncta per minute was then counted for the duration preceding administration and starting 10 min post administration.

#### Electron microscopy

Electron micrographs used in this study were collected for a previous study (19). Mouse hippocampi used for the postembedding immunogold localization were prepared as described previously (19–22). Mice were perfused with phosphate buffer, followed by perfusion with 4% paraformaldehyde + 0.5% glutaraldehyde in phosphate buffer. Fixed brains were vibratomed at 350 μm, then cryoprotected in glycerol overnight and frozen in a Leica EM CPC (Leica Microsystems, Wetzlar, Germany), and processed and embedded in Lowicryl HM-20 resin (Electron Microscopy Sciences) in a Leica AFS freeze-substitution instrument. Thin sections were incubated in 0.1% sodium borohydride + 50 mM glycine in Tris-buffered saline plus 0.1% Triton X-100 (TBST). Then they were immersed in 10% normal goat serum (NGS) in TBST, and primary antibody in 1% NGS/TBST (overnight), Then incubated with 10 nm immunogold-conjugated secondary antibodies (Ted Pella, Redding, CA) in 1% NGS in TBST with 0.5% polyethylene glycol (20,000 MW), and stained with uranyl acetate and lead citrate. In this material, the axonal compartment was identified definitively by its synaptic contact. In the original experiments for Sun et al., 2011 (19), a random sample of micrographs were taken from the hippocampus CA1-stratum radiatum from two experiments with 3+3 and 2+2 WT+KO mice; the two experiments produced similar results and the data were then combined for a total of 646 WT and 642 KO spine profiles. Only the WT spine synapse profiles from the 2011 study were used for this current study. The unpublished data from the analysis of the presynaptic terminals is presented here.

### Image analysis

#### Import and kymogram generation

Raw microscope time series were imported into ImageJ with StackReg and Bio-Formats plug-ins. A segmented line selection was drawn through the neurite of interest with thickness adjusted to cover the diameter of the dendrite (7-12 pixels), with a resultant ROI as shown in Figure S1A. Using the plugin KymoResliceWide, a kymogram was generated from the time series where the horizontal dimension corresponded to the average pixel intensity along the diameter of the cell for each pixel distance from the soma and the vertical dimension was time. A sample segment of kymogram from a bleached neurite is shown in Figure S1B. The intermittent photobleaching of the region of interest is marked on the sample kymogram with leftward facing blue arrows.

All kymograms were saved as TIF files, trajectories were saved as TXT files of coordinates, and all were imported into MATLAB for further processing.

#### Differentiating neurites

Axons and dendrites were differentiated based on their morphology. Dendrites exhibit a steady decrease in diameter with distance from the soma and terminate well within 1000 μm. Axons extend for thousands of microns and have a relatively constant diameter. The most obvious changes in diameter are in neurite trunks: the axon initial segment is thin like the axons, at a few microns, whereas dendritic trunks can be several microns thick and broadly blend into the plasma membrane of the soma. In addition, dendrites branch more frequently and at more acute angles, whereas axons can branch at perpendicular or even obtuse angles. Oftentimes, the morphological features differentiating axons and dendrites are not visible in the frame of the time series, and additional global images of neuron must be referenced to distinguish neurites. An example of this is depicted in Figure S1C, where the time series frame is outlined in red, but the defining morphological features of the axon and dendrite are only visible in the larger, global image. In Figure S1C, an axon (red arrow) and dendrite (blue arrow) exhibiting the aforementioned characteristics are labeled.

Beyond these morphological characteristics, the definitive way to differentiate neurites is with antibody staining for structural proteins exclusively found in one neurite type. Several coverslips were stained with ankyrin-G post live imaging to confirm identification of the axon initial segment. One such neuron is depicted in Figure S1D and Supplemental Video 2.

#### Contrast enhancement and thresholding

To improve the visibility of puncta trajectories, kymograms were enhanced using automated and manual methods in ImageJ. As an example, raw kymogram sections from a representative axon and dendrite are depicted in Figures S2A(i) and S2B(i). ImageJ’s automatic optimization of brightness and contrast is first performed based on the image’s histogram (Figures S2A(ii) and S2B(ii)). Next, the brightness and contrast settings were manually adjusted by narrowing the visible display range (Figures S2A(iii) and S2B(iii)). Lastly, a lower threshold was set, setting pixel values below this threshold to background, as shown in Figures S2A(iv) and S2B(iv).

#### Puncta trajectory selection

Puncta trajectories were traced using a segmented line selection. Oftentimes, the brightness/contrast and threshold settings were adjusted and readjusted for regions of varying immobile fraction within the same kymogram. For instance, a dendrite that is bleached five times over the course of a recording, as in Figure S1B, required different contrast and threshold settings to visualize puncta in early beaches and later bleaches.

In some cases, puncta appear to merge into one trajectory or split into multiple trajectories. An example of this is depicted in Figure S2C. In these cases, when tracing trajectories, each parent and child path is designated as an individual trajectory, as in the three trajectories depicted in Figure S2C(ii). The same protocol is followed for two puncta that seemingly merge into one trajectory.

Mobile puncta sometimes rapidly oscillate or vibrate in position. In these cases, if the specific path of the oscillations cannot be resolved, a trajectory was drawn through the mean position of the puncta. An example of this is depicted in Figure S2D, with a trajectory drawn through the mean position of an oscillation marked in Figure S2D(ii).

Further, a puncta can increase/decrease in fluorescence or appear/disappear during a recording, as shown in Figure S2E. Since segmented line selections are never drawn through neurite branch points, this likely corresponds to Kv4.2-SGFP2 dispersion or accumulation. To minimize subjectivity of trajectory selection through such events, each puncta trajectory was trimmed based on a threshold for net displacement, as described in the next section.

### Data analysis and modeling

#### Trajectory trimming

Since only mobile trajectories were considered, puncta with an immobile segment of trajectory before and/or after a mobile segment were trimmed. This was achieved by iterating through each trajectory and summing the net distance traveled. Portions of the trajectories up to the mobility threshold were removed, eliminating stall time before and after mobile segments. The minimum distance threshold was 5 μm for both axon and dendrite trajectories. As an example, both trajectories shown in Figures S2E(ii) and S2E(iii) are interpreted as the same trajectory (Figure S2E(iv)) following trimming. This was useful in cases where puncta appear or disappear on a kymogram, as in Figure S2E(i). This also relieves some degree of subjectivity surrounding puncta start/end points and in measurement of stall time.

#### Total distance traveled vs net displacement

The total distance each Kv4.2-SGFP2 puncta travels was computed by summing the absolute value of the distance traveled between each frame in a time series. Moreover, the net displacement equals the puncta final position minus initial position.

#### Average speed and stall time

To average speed, the instantaneous velocity was computed between each two frames of a time series. The mean of the absolute values of these instantaneous velocities equals the average speed. Puncta stall time is defined as the fraction of total time during which puncta are traveling with a speed less than 0.1 μm/sec.

#### Mean squared displacement and super-diffusion

Mean squared displacement (MSD) was computed by averaging the square of the difference between puncta coordinates some time τ apart. This was repeated for τ up to one fourth the length of the recording duration. MSD was then plotted against τ, and equation MSD(τ) = *D*τ^*α*^ was fit for each set of coordinates. *D* and α were then recorded as diffusion and superdiffusivity coefficients, respectively.

#### Steady state analysis for compartmental model

The cargo content of each compartment in a model is defined by a differential equation that sum the quantities of cargo entering and exiting that compartment. A generalized rate *v*_*d,r*_ from a donor *d* to receiver *r* transfers an amount of mass *dv*_*d,r*_. As an example, the system of differential equations for the simplest model in Figure S6B is as follows:

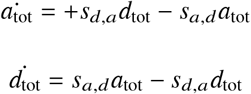

This system of equations can be solved at steady state to estimate the ratio of these rates. The steady state assumption sets 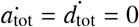. Then, rearranging either equation yields

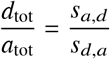

We can now use our experimental data to restrict rates between axon and dendrite.

#### Stochastic model fit to experimental data

To fit the stochastic model of a random walk modified with offload rate (*p*_off_) and memory (*p*_mem_) to experimental data, we use a combination of maximum likelihood estimation (MLE) and least squares fitting. Experiment and model data are normalized and fit to a gamma distribution using MLE (MATLAB function fitdist). A gamma distribution accommodates all data given its continuity and coverage of a semi-infinite [0,∞) interval.

The shape and scale parameters of gamma fits are compared using nonlinear least squares data fitting (MATLAB function `lsqcurvefit`). Generating stochastic model estimates requires a large number of simulated puncta *N*_*s*_ to produce consistent distributions. To resolve this, we employ a moderate *N*_*s*_ = 10, 000 and increase the finite difference step size of `lsqcurvefit`. A script continuously iterates between (1) running N iterations of the stochastic model, (2) MLE of stochastic data, and (3) least squares fitting of distribution parameters to match those of experimental data. A full description of this heuristic method is presented in Supplementary Materials.

## RESULTS

### Mass action transport causes discordance in cargo densities between microtubule and surface compartments

The question we address in this study is how densities of actively transported cargo relate to membrane-bound localization. We begin by discussing a conceptual model of how cargo densities on microtubules give rise to functional densities in the plasma membrane. A cartoon of the potential relationships is shown in Figure 1. Cargo on microtubules and cargo localized to the plasma membrane is depicted by the shading of the interior and outline of the cell, respectively (Figure 1A). Intuitively, one might expect to measure higher densities of transported cargo in the compartments where the cargo eventually becomes localized (Figure 1B). Indeed, this is the implicit assumption made in static imaging studies that attempt to quantify intracellular protein and mRNA distributions by labeling and counting puncta or integrating signal density (23).

**Figure 1:**
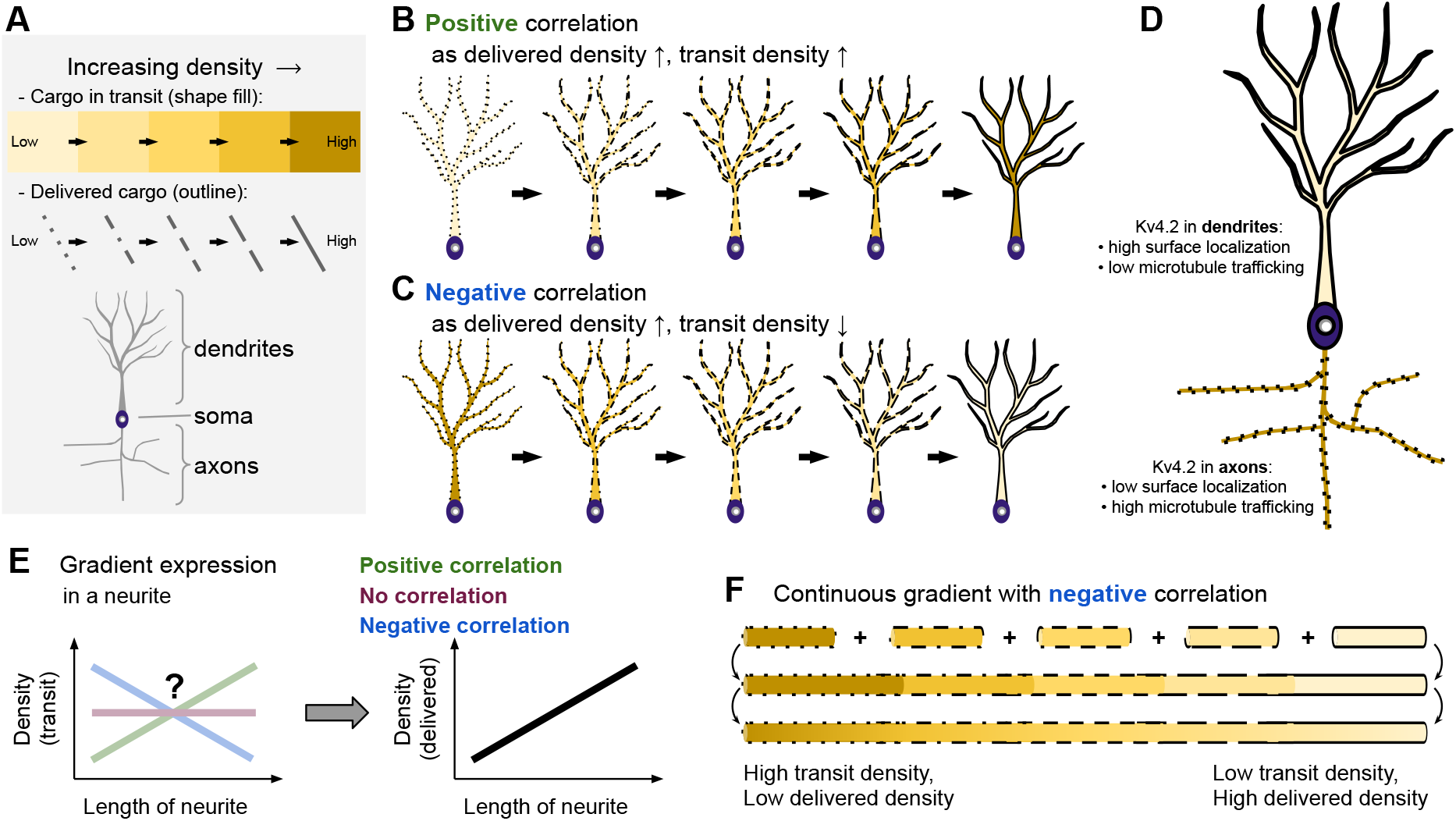
Mass action transport causes discordance between delivered cargo densities cargo in transit. (A): Schematic legend. (B): If membrane and microtubule densities are positively correlated, both quantities increase or decrease together. (C): In a negative correlation, an increase or decrease in membrane density results in the opposite deflection in the microtubules. (D): Measured densities for Kv4.2 are negatively correlated, consistent with mass action models for a system with multiple neurites. (E): This principle can be applied to gradient expression along a single neurite. The question of positive, negative, or no correlation remains. (F): Kv4.2 exhibits continuous gradients of cargo delivered and in transit - consistent with negative correlation.

However, under mass-action, the rate of transport between cellular compartments is proportional to density. In a well mixed system with homogeneous cargo affinity, cargo with strong membrane affinity can fill both microtubule and membrane compartments simultaneously, as in Figure 1B. With heterogeneous, compartmentalized neurites, the microtubules in branches with lower membrane affinity act as a cargo sink. This can result in negative correlation between surface expression and membrane expression within each section of neurite, as depicted in Figure 1C.

We experimentally measured and quantified the microtubule trafficking of a specific cargo, Kv4.2 subunits, whose distribution is especially relevant to these considerations. Kv4.2 has a highly regulated dendritic surface expression whose density increases along dendrites with distance from the soma. We measured membrane density and microtubule density of Kv4.2 in neurites of varying surface expression. A summary of these results is depicted in Figure 1D. Kv4.2 has high surface localization and surprisingly low microtubule density in dendrites. We show that a gradient density of delivered cargo can have a positive, negative, or no correlation with the gradient of that cargo in transit (Figure 1E). We find that the density of Kv4.2 subunits in transit decreases in dendrites with distance from the soma. Taken with localization data and the well-established functional profile of the channel, this is consistent with a continuous expression gradient with negative correlation, depicted in Figure 1F.

### Kv4.2 microtubule-based trafficking is observed more frequently in axons than in dendrites

To establish reliable estimates of the frequency, density, and kinetic properties of actively transported Kv4.2 subunits we performed 129 hour-long recordings neurites in cultured rat hippocampal cells. In total 507 mobile Kv4.2-SGFP2 puncta were identified among 478 recorded dendrites, and 961 mobile puncta were identified in 46 axons (see Methods). The durations over which puncta are mobile are depicted in Figure 2A for axons and Figure 2B for dendrites. Of the 478 dendrites in hour-long recordings, only 213 dendrites (45%) exhibited at least one mobile punctum. Only data from this subset of dendrites is presented in Figure 2B. Mobile puncta appeared consistently in axons, whereas in dendrites mobile puncta appear intermittently or not at all. The average length of a sampled region was 85.4 μm in axons compared to 52.3 μm in dendrites. We found no strong correlation between transit frequency and degree of branching, from primary (apical) dendrites to quaternary branches (Figure S3). To ensure puncta visibility was not an artifact of fluorescence intensity, we plot puncta frequency versus standardized neurite intensity and find no strong correlation (Figure S4). We do find a correlation between puncta frequency and distance from soma in dendrites (Figure 2F), which is used for analysis in subsequent results (Figure 5).

**Figure 2:**
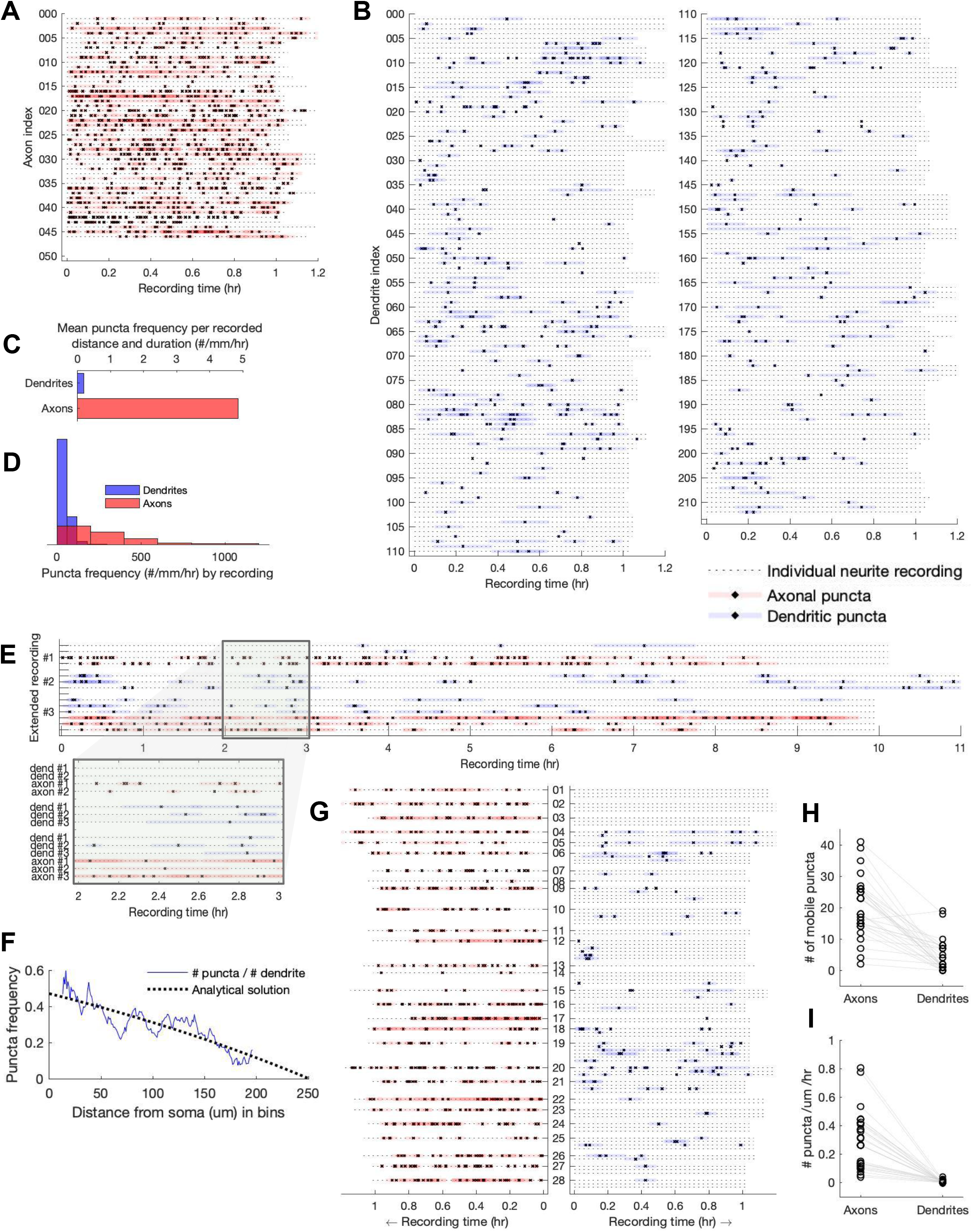
Kv4.2 microtubule-based trafficking is observed more frequently in axons than in dendrites. (A): Hour-long recordings of 46 axons are depicted, with highlighted sections indicating periods of puncta mobility. (B): Hour-long recordings of 213 dendrites are depicted, with highlighted sections indicating periods of puncta mobility. This subset of 478 dendrites has ≥ 1 mobile puncta. (C): Puncta frequency in axons and dendrites is standardized by total neurite length visualized and time recorded (units: number of puncta/mm/hr). (D): Histogram depicting puncta frequency by neurite recording. (E): Three extended recordings that substantiate the puncta frequency discrepancy between axons and dendrites over extended periods of observation. (F): Puncta frequency decreases with distance from soma in dendrites, consistent with analytical solutions to the drift-diffusion equation. (G): Axons and dendrites originating from the same soma (same neuron) are depicted, demonstrating similar trends as those observed in isolated recordings. The central column of numbers indicate an arbitrary recording index. (H): Number of mobile puncta per neurite for concurrent recordings from (G). (I): Number of mobile puncta per neurite standardized by length and time for concurrent recordings from (G).

When standardizing these measurements for recording duration and neurite length, the discrepancy in mobile puncta frequency is 4.9 puncta/mm/hr in axons versus 0.18 puncta/mm/hr in dendrites, depicted in Figure 2C. Puncta frequency in dendrites drops to 0.039 puncta/mm/hr when considering dendritic recordings with zero mobile puncta (not depicted). A histogram showing puncta frequency by neurite recording is depicted in Figure 2D.

In order to control for the possibility of global trafficking failure in dendrites that did not show puncta during hour-long recordings, we performed extended recordings lasting 10 hours, shown in Figure 2E. The trend in puncta frequency for extended recordings is consistent with that of hour-long recordings, suggesting that hour-long recordings with no puncta are simply a result of sampling.

In some cases, it was possible to reliably identify and record from axons and dendrites originating from the same soma to control for intercellular variations in trafficking or metabolism. Axons and dendrites from these 28 recordings are depicted alongside each other in Figure 2G. In all but one case, axons possessed the majority of mobile puncta, even though multiple dendrites were recorded for most neurons. Comparisons of raw puncta count (Figure 2H) and standardized puncta count (number of puncta/μm/hr) (Figure 2I) are depicted. After standardizing measurements to sampling distance and duration, the axons average a 36-fold increase over the simultaneously recorded dendrites from the same cell.

To validate that mobile puncta are transported via active, motor protein based transport we applied the microtubule-disrupting drug colchicine (24–26). Six coverslips were treated with colchicine during hour-long recordings. On average, colchicine administration resulted in a substantial (> 60%) decrease in number of mobile puncta in axons when compared to vehicle (Figure S5). The Kv4.2-SGFP2 puncta transport that we observe elsewhere is thus likely to be an active, microtubule dependent process.

Thus far, we have imaged and analyzed mobile fractions of Kv4.2. Together, these data establish that actively transported Kv4.2 puncta are present in significantly higher densities in axons as compared to dendrites. Next we measured the static and endogenous Kv4.2 density in dendrites and axons.

### Kv4.2 preferentially localizes to plasma membrane of dendrites compared to axons in both endogenous and transfected expression systems, but expression in axons is not negligible

To independently assess the static density of native Kv4.2 expression in axons and dendrites, we used electron microscopy (EM) following immunogold labeling of endogenous Kv4.2 subunits. Owing to the inherent constraints of EM imaging, we quantified axon/dendritic expression in identifiable pre- or postsynaptic compartments, respectively, and regions of neurite that were clearly contiguous with these compartments. We imaged 624 presynaptic and 646 postsynaptic regions. Example micrographs in Figure 3A show an axon (ax) that can be traced to presynapses (pre), with gold particles present in both the axon shaft and terminals.

**Figure 3:**
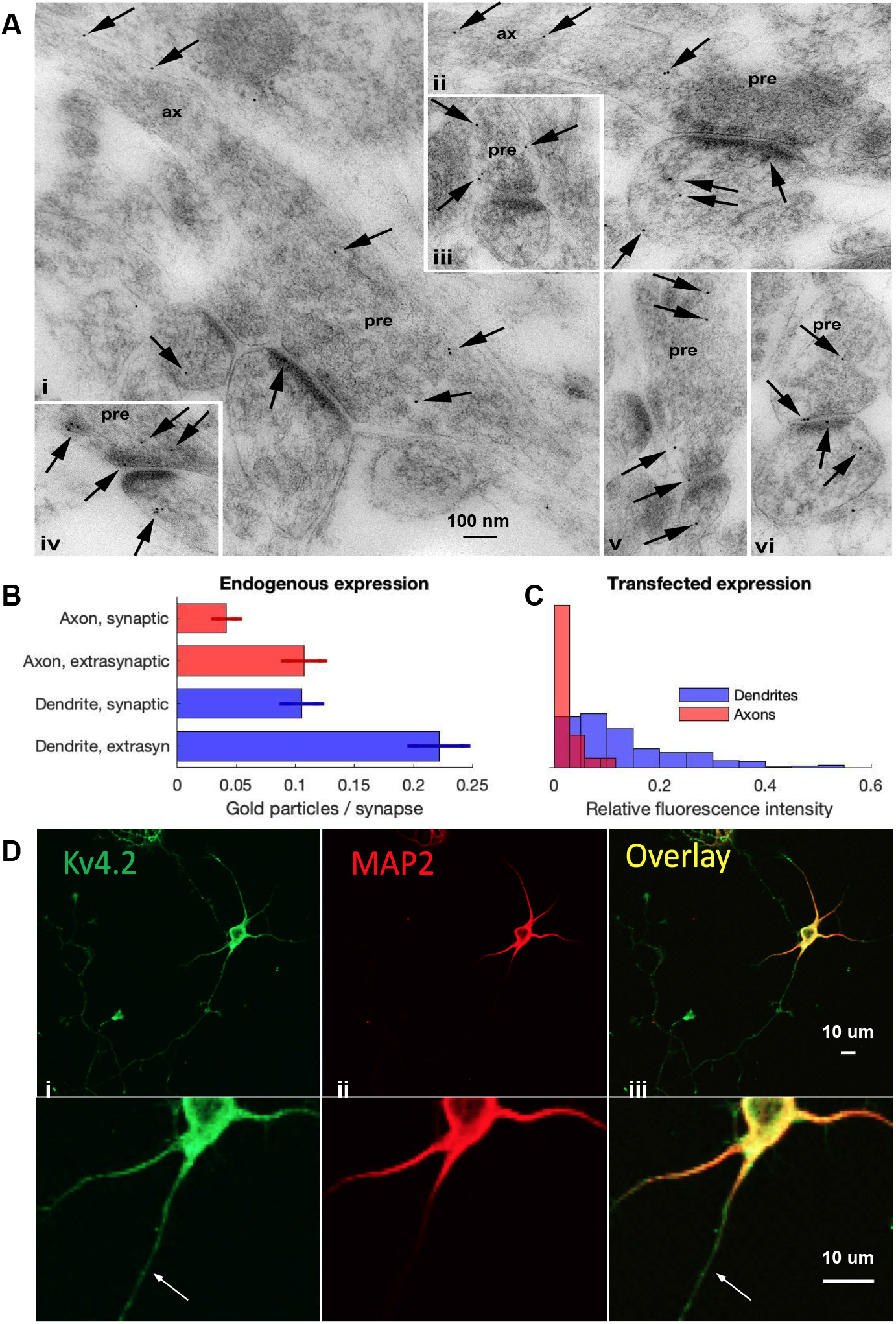
Kv4.2 preferentially localizes to plasma membrane of dendrites compared to axons in both endogenous and transfected expression systems, but expression in axons is not negligible. (A): Immunogold localization (arrows) of Kv4.2 in the CA1 stratum radiatum of the hippocampus of WT mice. Synapse profiles show the presynaptic terminal (pre) contacting one or two postsynaptic spines. In (i) and (ii), the axon (ax) can be traced from the presynaptic terminal. Gold labeling extends along the axon and into the presynaptic terminals. Examples of gold labeling associated with the plasma membrane of the synapse and counted in the accompanying graph include those at the axon synaptic membrane shown in (iv), (v), and (vi), the axon extrasynaptic membrane shown in (iii) and (v), the dendrite synaptic membrane shown in (i) and (vi), and the dendrite extrasynaptic membrane shown in (ii). (B): Quantification of (A). Concentrations in the four compartments are 0.0417, 0.1074, 0.1053, and 0.2214 gold particles per synapse. Error bars indicate standard error of the mean. Note that the bars for ‘Dendrite, synaptic’ and ‘Dendrite, extrasyn’ were published previously in another form in Sun et al., 2011 (19) and are represented here for comparison. (C): Histogram of the relative prebleach fluorescence intensity for axons versus dendrites from Kv4.2-SGFP2 transfected dataset, consistent with electron microscopy here and in previous studies. (D): E18 cultured rat hippocampal neurons at DIV5 were immunostained with Kv4.2 ((i), green) to visualize the endogenous Kv4.2 and MAP2 ((ii), red) to mark the dendritic arbor. The arrow indicates an example of an axon that still shows substantial endogenous Kv4.2.

Sampled immunogold particles identified in the synapses and perisynapses are broadly divided into pre/postsynaptic regions. Presynaptic terminals (axons) contained 30.6% of all gold particles and 0.15 particles/synapse. Postsynaptic terminals (dendrites) contained 69.4% of particles and 0.33 particles/synapse. This is consistent with previous localization studies (8) in showing substantial, non-negligible subunit localization in axons. The pre/postsynaptic regions are subdivided into synaptic and extrasynaptic regions. In both axons and dendrites, under one third of particles (28.0 and 32.2%, respectively) of particles were found in synaptic spaces, with the remaining two thirds in extrasynaptic regions. These percentages and gold particle frequencies are summarized in Table 1 and depicted in Figure 3B.

**Table 1:** Density of immunogold particles identified by electron microscopy in synapses of axons and dendrites.

| Axons, N = 624 |  | Dendrites, N = 646 |  | Neurite, number of synapses sampled |
| --- | --- | --- | --- | --- |
| 93 |  | 211 |  | Number of gold particles |
| 0.149 |  | 0.327 |  | Gold particles / synapse |
| 30.6 |  | 69.4 |  | Percent of total |
| *Synaptic* | *Extrasynaptic* | *Synaptic* | *Extrasynaptic* | Subdivision |
| 26 | 67 | 68 | 143 | Number of gold particles |
| 0.042 | 0.107 | 0.105 | 0.221 | Gold particles / synapse |
| 28.0 | 72.0 | 32.2 | 67.8 | Percent of subdivision |
| 8.6 | 22.0 | 22.4 | 47.0 | Percent of total |

Immunolabeling further confirmed endogenous localization of Kv4.2 in axons. Figure 3D depicts a neuron with multiple dendrites and one axon stained for somatodendritic marker MAP2 ((ii), red). Neurites lacking this marker are designated as axons. We found substantial Kv4.2 ((i), green) in both dendrites and the axon (marked with the arrow). Kv4.2 observed in the axon is well above levels of background staining, providing further evidence of non-negligible subunit density in axons. Our measurements of Kv4.2-SGFP2 transfected neurons also corroborates this trend. A histogram of the prebleach fluorescence intensity is depicted in Figure 3C. Dendrites appear to contain significantly more Kv4.2 per unit area compared to axons. Taken together, these results establish that Kv4.2 preferentially localizes in dendrites, but its expression in axons is non-negligible, consistent with other studies.

### Disparity between Kv4.2 static localization and mobile frequency is explained by a mass-action model

Our results so far show that the static distribution of Kv4.2 is concentrated in dendrites (Figure 3), in agreement with previous studies and known physiological function of this channel. On the other hand, we find that the vast majority of subunits undergoing active transport appear in axons (Figure 2). How can these apparently conflicting results be reconciled? We addressed this question by constructing biophysical models of active transport that we constrained with the transport frequencies we obtained from experiments. Such models enable us to infer whether the disparity in actively trafficked axonal and dendritic subunits (Figure 2) is consistent with the opposite density in localization (Figure 3).

A full neuron morphology can be discretized into spatial compartments, depicted in Figure 4Ai. In each compartment, we assumed that cargo is either undergoing transport on microtubules (subscript *mt*) or delivered (subscript *del*) in axonal (*a*) and dendritic (*d*) compartments. Compartments *del* account for all channel subunits that have detached from microtubules, including those in local pools and on the plasma membrane. Rates from *mt* to *del* represent cargo offloading from the microtubules (*a*_off_, *d*_off_), lumping together microtubule release, actin filament loading/unloading, local trafficking, and any local modification before insertion of the channel into the plasma membrane. The reverse rates (*a*_reload_, *d*_reload_) represent cargo reloading from *del* to *mt*. Our puncta measurements sampled segments of dendrites and axons, which did not provide data at all locations along each neurite. To incorporate these measurements into a model, we coarsened into a lumped compartmental model that considers only the average density of material in axons and dendrites, irrespective of location (Figure 4Aii). In the lumped model, *s*_*a,d*_ and *s*_*d,a*_ represent cargo passing between axons and dendrites. All other rates and compartments are as previously described.

**Figure 4:**
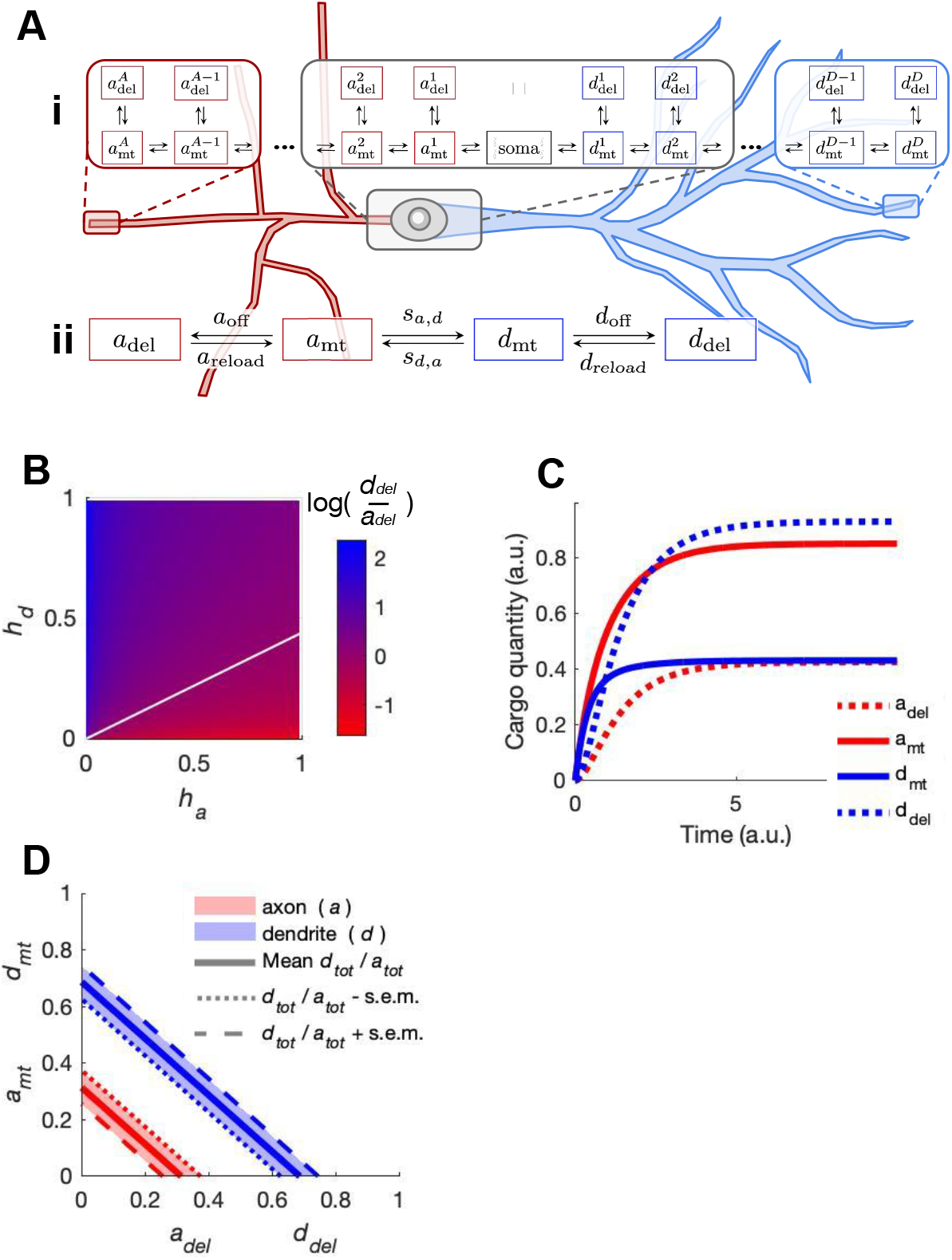
Disparity between Kv4.2 localization and mobile frequency is explained by mass-action transport kinetics. (A): Box diagram of mass action model of axon and dendrite transport. In a full morphology (i), the central soma is surrounded by microtubule (*mt*) and delivered (*del*) cargo compartments for axons *a* and dendrites *d*. Arrows denote rates of cargo transfer between compartments. A lumped variant (ii) can accommodate experimental constraints to simulate disparities in subunit density between axons and dendrites. (B): Estimated ratio *d*_del_/*a*_del_ as a function of delivered fractions *h*_*d*_ and *h*_*a*_. The white line indicates *d*_del_/*a*_del_ = 1. (C): Result of simulation with experimentally-constrained rates, corroborating transit frequency discrepancy observed experimentally. (D): Analytical result demonstrating negative correlation between *a*_del_ and *a*_mt_ or *d*_del_ and *d*_mt_ when restricted to a constant *tot* mass.

The system of differential equations for this lumped compartmental model is show below:

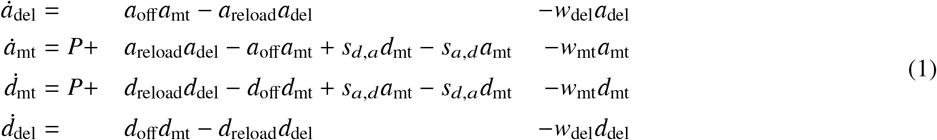

where a lumped rate *v*_*d,r*_ describes mass flow from state *d* to state *r*. The lumped model does not contain a soma compartment. To account for biosynthesis, we add a fixed production term *P* to both dendritic and axonal microtubule compartments. Note that flux into both is not assumed to be equal, since flow between axons and dendrites is accounted for by parameters *s*_*d,a*_ and *s*_*a,d*_. *w* represents cargo degradation, which, consistent with endolysosomal and authophagic degradation pathways of membrane proteins (27), is faster in *del* than *mt*: *w*_del_ > *w*_mt_. The remaining rates in Eqs. 1 are estimated from experimental results.

We assumed that *s*_*a,d*_ and *s*_*d,a*_ are slower than the other four rates for two reasons. First, the distances traveled on microtubules are substantially longer than from *mt* to *del*, as evident in the full morphology in Figure 4Ai. The net flux due to active transport between dendritic and axonal compartments is lumped into the parameters *s*_*a,d*_ and *s*_*d,a*_. Allowing separate fluxes, *s*_*a,d*_ and *s*_*d,a*_, provides for asymmetric flow due to sorting mechanisms that are known to regulate cargo entry into both axons and dendrites (28–30), including for Kv4.2 (31). To enable a (quasi) steady state estimate of cargo density we set rates *s*_*a,d*_ and *s*_*d,a*_ to a timescale ten-fold slower than the other rates, although more modest timescale separation produced the same qualitative result.

We next constrained the rates in this model with our experimental measurements. Rates *s*_*a,d*_ and *s*_*d,a*_ are estimated using the total cargo in axons *a*_tot_ and dendrites *d*_tot_. We assumed that

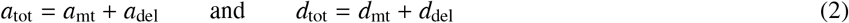

and that immunogold labeling of Kv4.2 captures *a*_tot_ and *d*_tot_. We found that roughly two-thirds of subunits were present in dendrites and one-third in axons (see Figure 3B and Table 1). Predominant dendritic segregation of the channel is corroborated by other localization studies (4, 6, 8, 11) as well as our measure of transfected fluorescence intensity per unit area (Figure 3C). To estimate rates from *tot* densities, we grouped *mt* and *del* compartments (from Figure 4Aii) to produce a model with only *a*_tot_ and *d*_tot_, depicted in Figure S6B. Steady state analysis of the differential equations (see Methods) for this simplest model gives:

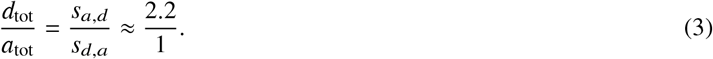

where we approximated *s*_*a,d*_/*s*_*d,a*_ using *tot* densities from Figure 3B and Table 1, normalized to axonal measures.

Estimating offload (*a*_off_, *d*_off_) and reload (*a*_reload_, *d*_reload_) rates requires a measure of *mt* and *del* cargo in both axons and dendrites. We constrained the steady state densities of *mt* compartments (*a*_mt_, *d*_mt_) using experimental data (Figure 2C). Normalizing to axonal puncta frequency, we estimated *d*_mt_ = 0.04 and *a*_mt_ = 1.

Inferring *a*_del_ and *d*_del_ is more difficult. We have no raw data of surface-expressed Kv4.2, since it cannot be distinguished from subunits in transit. A-type current through Kv4.2 has not been reported in axons, so electrical or functional data cannot be used. We cannot compute *del* using Eq. 2, since *tot* and *mt* have nonconvertible units. We therefore proceeded by parameterizing *del* in terms of *tot* using *a*_del_ = *h*_*a*_*a*_tot_ and *d*_del_ = *h*_*d*_*d*_tot_. In this notation, *h*_*a*_ and *h*_*d*_ are the fractions of delivered cargo in each neurite.

These manipulations allowed us to sweep across all plausible relative densities of surface-expressed Kv4.2, given the measurements that we could obtain from data. Figure 4B shows all possible *d*_del_/*a*_del_, where the white line indicates *d*_del_ = *a*_del_. From Figure 4B, *d*_del_ < *a*_del_ only occurs for *h_d_* < 0.43 and large *h*_*a*_, both of which are unlikely for the following reasons. Dendrites have low subunit trafficking frequency and high prebleach intensity (see Figures 2D and 3C). Axons likely have low *h*_*a*_ given their low prebleach intensity (Figure 3C) and no detectable (reported) Kv4.2 current. It is therefore probable that *d*_del_ > *a*_del_. For our simulation, we assume that *h*_*a*_ = *h*_*d*_ for the conservative estimate that *d*_del_/*a*_del_ = 2.2. Normalizing to the axon, we use *d*_del_ = 2.2 and *a*_del_ = 1.

To estimate offload and reload rates from *mt* and *del* densities, we modeled axons and dendrites individually as depicted in Figure S6C. As before, we arrived at expressions that allowed us to solve for ratios of transport/surface expression rates:

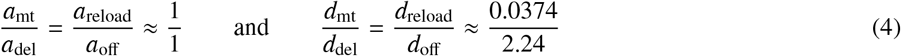

Together these estimates provide constraints for all rates in the lumped model (Figure 4Aii).

The solutions of this model are depicted in Figure 4C. The negative correlation between *mt* and *del* compartments of both neurites is clear: *a*_del_ < *a*_mt_ and *d*_mt_ < *d*_del_. In the context of mass action, the result is intuitive. Because surface localization in axons is restricted (*a*_del_ < *d*_del_), more cargo tends to accumulate in the microtubules of axons versus those of dendrites (*a*_mt_ > *d*_mt_). The apparent paradox of increased Kv4.2 trafficking in regions of low Kv4.2 function/expression can therefore be explained by a simple mass action model constrained by measurements of trafficking kinetics.

We examined the robustness of a negative correlation between *mt* and *del* compartments using Eq. 2 and the estimate in Eq. 3. We normalized the constraint in Eq. 3 to a total mass *a*_tot_ + *d*_tot_ = 1 such that each variable is a fractional quantity: *a*_tot_ = 0.31, *d*_tot_ = 0.69. The resulting steady-state densities of delivered and transported cargo is plotted in Figure 4D. Shaded regions indicate the range of *a*_tot_ *d*_tot_ over one s.e.m. using data from Figure 3B. Quantities of cargo *mt* have a clear negative correlation with *del*. This relationship holds for any *a*_tot_ *d*_tot_.

In summary, these results so far show that apparently contradictory densities of mobile and surface-bound cargo are consistent with a simple transport model. This conclusion only required considering bulk flow between compartments that represented the entirety of the axonal and dendritic arbors. However, our experimental measurements also indicated strong differences in the detailed motion of axonal and dendritic puncta. We next analysed detailed transport kinetics in both axons and dendrites to infer whether differences could be accounted for by qualitatively different transport mechanisms.

### Kinetic differences between axons and dendrites are attributable to varying propensities for cargo offloading and unidirectional runs

We analysed Kv4.2 puncta trajectories in axons and dendrites. Kymograms depicting representative puncta trajectories in axons and dendrites are shown in Figure S7A(i) and Figure S7B(i), respectively. Population measurements of puncta kinetics are shown in Figure 5. We found that axonal puncta undergo unidirectional runs at high speeds, whereas dendritic puncta appear to change direction more frequently and stall longer. Since the same cargo is trafficked in both neurite types, we asked whether these differences in trafficking kinetics were better explained by an intrinsic difference in the active transport mechanism, or whether they could simply be accounted for by a difference in dendritic and axonal propensities for cargo being offloaded to the membrane. To infer which of these mechanisms best captured our observations we modelled out data using a modified random walk.

**Figure 5:**
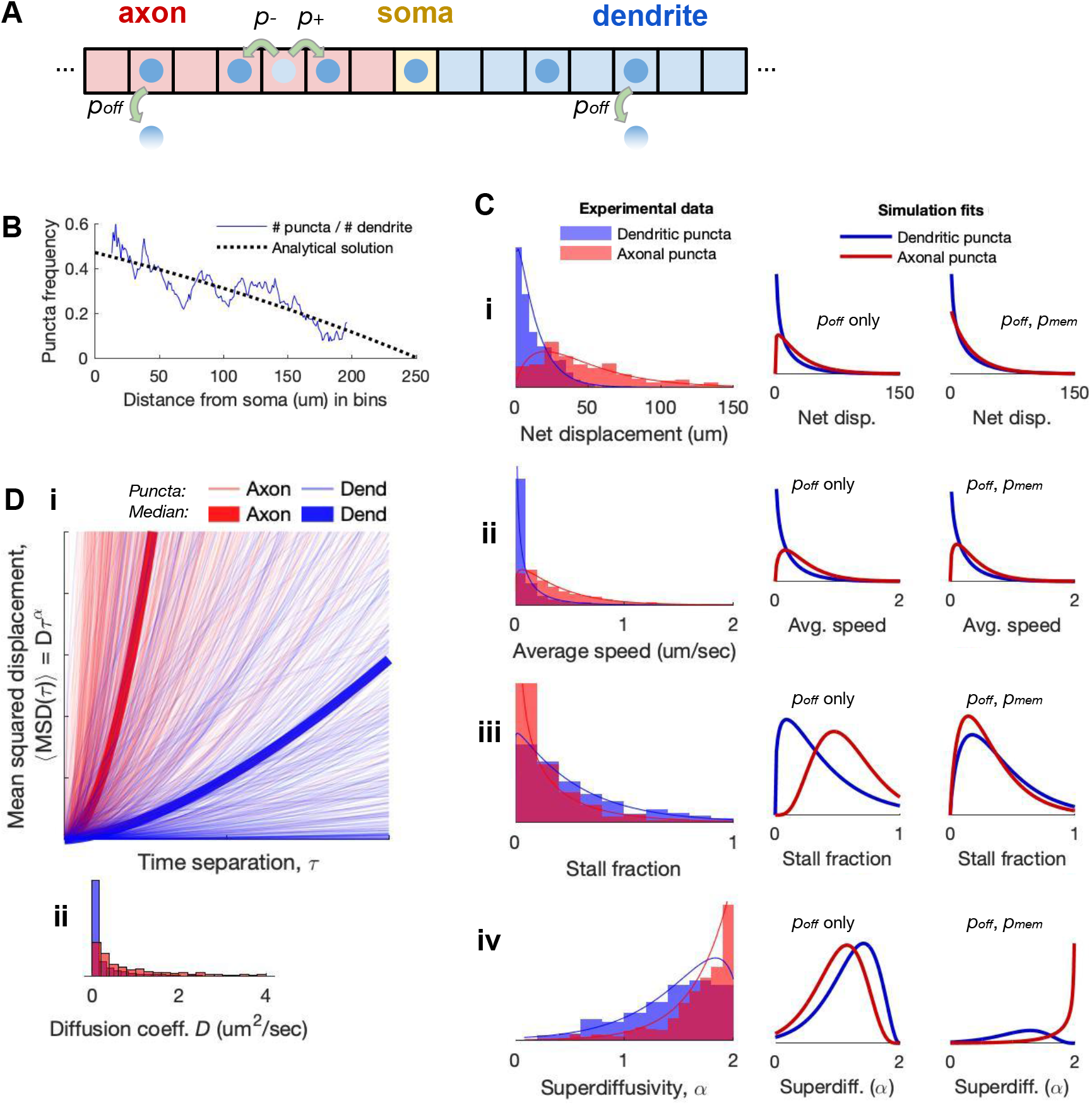
Kinetic differences between axons and dendrites are attributable to varying propensities for cargo offloading and unidirectional runs. (A): Setup of stochastic simulations along linear multi-compartment model (axon-soma-dendrite), with left/right jump and offloading rates depicted. Complete model is depicted in Figure S8A. (B): Puncta frequency decreases with distance from soma in dendrites, consistent with analytical solutions to the drift-diffusion equation. (C): Histograms for various transport parameters, normalized as probability density functions. On average, axonal puncta have greater net displacement (i), faster speed (ii), deceased stall time (iii), and increased unidirectional runs (iv). Model fits to these results using *p*_off_ alone (*second column*) as well as using *p*_off_ and *p*_mem_ (*third column*) are depicted. (D): Result of curve fitting for mean squared displacement (MSD) versus time separation (*τ*), revealing higher degree of superdiffusivity in axons compared to dendrites (i). Bold curves indicate the medians of the populations. Histogram show distribution for *D* (ii) and *α* (Civ).

#### modeling transport using a modified random walk

We constructed a simplified compartmental model of active transport in the form of puncta undergoing modified random walks. This model is shown in Figure 5A and contains three types of compartment: axon (A), soma (S), and dendrite (D). Each punctum begins in the S compartment and has some probability per unit time (propensity) of moving right *p*_+_ and some propensity of moving left *p*_−_, such that *p*_+_ + *p*_−_ = 1. These transition propensities represent the stochastic movement of actively transported cargo along microtubules.

We incorporated surface expression dynamics by allowing surface compartments, with neurite-specific propensities for trafficking between the microtubule and surface. Puncta in axons have a propensity of surface expression, 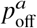, while puncta in dendrites have a distinct propensity 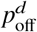. Given our own observations (Figure 3) and the extensive published evidence for Kv4.2 proclivity to dendritic versus axonal surface expression (4, 6–8, 11, 14, 32), we explored ratios of 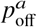 to 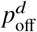 for which 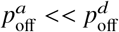.

In a variation of this model, we implemented our observation that axonal puncta travel unidirectionally with little stall time, which is characteristic of a superdiffusive process. Indeed, there is experimental evidence for microtubule-based transport undergoing superdiffusive unidirectional runs (33–35). We therefore incorporated an additional probability parameter *p*_mem_ that acts as a memory term, depicted in Figure S8Aii and iii. *p*_mem_ is the probability that a punctum repeats its previous step, giving rise to extended runs if *p*_mem_ > 0.

Figure S8B depicts a simulation of 10 puncta over 500 time steps. The resultant simulated trajectories agree qualitatively with kymograms from experiments (Figure S7A-B). In this simulation, as in experiments, puncta in axons on average travel longer distances, at faster speeds, and with decreased stall times when compared to puncta in dendrites. We next rigorously examined the goodness of fit of our experimental data to this model to infer any mechanistic differences between axons and dendrites.

We estimated left and right jump rates *p*_−_ and *p*_+_ of individual puncta by analyzing the bulk flow of a population of particles. We used population dynamics for average puncta position as a function of distance along the dendrite. Figure 5B depicts the observed puncta frequency of dendritic puncta versus their measured distance from the soma. Puncta trajectories are grouped in 1 μm bins along the dendrites, normalized by the number of dendritic recordings in each bin. To avoid numerical errors with low replicate count, we only considered bins with ≥ 30 dendrite recordings. The resulting distribution of puncta frequency is plotted (Figure 5B) and displays a trend of decreasing puncta frequency with distance from the soma.

This distribution of puncta frequency versus distance is expected for a collection of mobile particles obeying the drift-diffusion equation, which we can demonstrate analytically. The one-dimensional drift-diffusion equation is as follows:

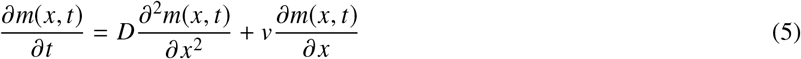

where *m(x,t)* denotes the concentration of some substance (heat, particles—in this case, Kv4.2-containing puncta) as a function of position *x* and time *t*. *D* is the diffusion coefficient and *v* is the mean net velocity, or drift.

Since all of our measurements were in cells with strong fluorescence many hours after transfection, we may assume transport has reached a steady state in which an equal number of puncta enter and leave the recording region. Thus, 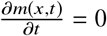, reducing Eq. 5 to:

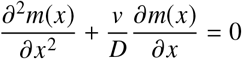

This special case of the drift-diffusion equation is known as Poisson’s equation, which we can solve as a boundary value problem using the boundary conditions observed experimentally. The endpoints of our data, *m*(0 μm) = *B*_*P*_ and *m*(200 μm) = *B*_*D*_, are set for fitting, where *B*_*P*_ and *B*_*D*_ are also the proximal and distal boundaries of the model. Our analytical solution is as follows:

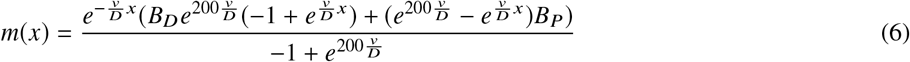

We fitted this analytical solution to the experimental data using least squares to obtain *B*_*P*_, *B*_*D*_, and the drift/diffusion coefficient ratio 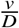:

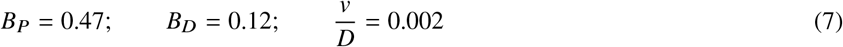

This analytical solution is overlaid on the experimental data in Figure 5B.

*D* and *v* describe the bulk flow of a population of particles. When Eq. 5 is discretized, *D* and *v* characterize the rates of cargo transfer between adjacent compartments:

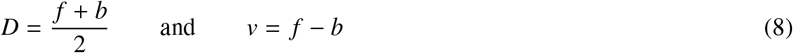

where *f* and *b* are the forward and backward rates of the discretized compartmental model. In the limit of large numbers, the propensities of a particle undergoing a random walk *p*_+_ and *p*_−_ are related to compartmental model rates *f* and *b* according to

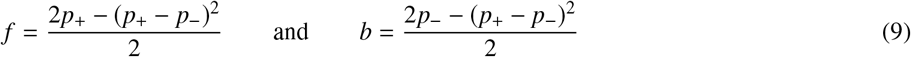

as derived in (5). Using the result from the BVP (Eq. 7) along with Eqs. 8 and 9 and the constraint that *p*_+_ + *p*_−_ = 1, we can estimate random walk jump rates: *p*_+_ = 0.5005, *p*_−_ = 0.4995. Puncta thus have a modest directional bias since *p*_+_ ≈ *p*_−_.

### Inference of qualitative differences in axonal and dendritic trafficking via stochastic modeling of experimental data

Returning to the random walk model, we performed parameter inference of model parameters using experimental data and maximum likelihood (see Methods and Supplemental Materials). We compared models with (0 < *p*_mem_ < 1) and without (*p*_mem_ = 0).

Figure 5C (second column) shows fits to a memoryless model (*p*_mem_ = 0). Optimal parameter estimates for surface delivery gave 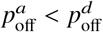 (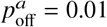 and 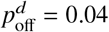) consistent with experiments and previous results in Figure 4. A random walk with *p*_off_ alone is sufficient to explain the differences in net displacement and average speed (Figure 5Ci and ii). However, stall time and unidirectional runs are not captured (Figure 5Ciii and iv).

The result of fitting the model with a memory term (0 < *p*_mem_ < 1) is shown in the third column of Figure 5C. We again found 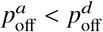, producing the same trends in displacement and speed (Figure 5Ci and 5Cii). Optimal estimates of the memory term were 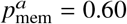 and 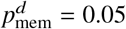 which is consistent with superdiffusivity being prominent in axons (Figure 5Ciii and 5Civ) and with longer unidirectional runs and with elevated stall times in dendrites, as observed in trajectories (Figure S6A-B). These observations are consistent with an analysis of mean squared displacement (MSD) for each trajectory (Figure 5D). We computed average MSD as a function of time interval, τ up to one quater of the total recording duration. Resulting data were fitted to MSD(τ) = *D*τ^*α*^ for each trajectory to obtain parameters *D* and α. Puncta in both dendrites and axons appear to undergo motion with similar *D* (Figure 5Dii). However, the MSD tends to increase more rapidly with τ for axonal puncta than for dendritic puncta (Figure 5Di). This corresponds to the axonal puncta taking more consecutive steps in the same direction, resulting in motion that is more directed than the memoryless walk of particles in typical diffusion. Axon puncta exhibited a higher degree of superdiffusivity than dendrite puncta (Figure 5Civ). This discrepancy is consistent with simulated puncta behavior with inferred parameters 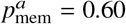 and 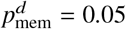.

Together, this analysis suggests mechanistic differences in the transport of Kv4.2 in axons and dendrites. Increased net displacement, average speed, and puncta frequency in axons are explained by a random walk with minimal delivery in axons, consistent with Kv4.2 predilection for dendrites. However, reduced surface delivery of cargo in axons only partially explains the longer observed runs. Other kinetic parameters - stall time and superdiffusivity - require an additional memory term in a modified random walk model, suggesting a distinct axonal transport mechanism.

### Opposing gradients in static localization and mobile frequency along somatodendritic axis are reconciled with mass-action kinetics

We have established that lumped densities of delivered cargo (*del*) and cargo in transit (*mt*) are negatively correlated (Figures 2–3), consistent with models of mass action (Figure 4). In distance-dependent measurements, we observe a decreasing *mt* profile along dendrites with distance from the soma (Figure 5B). On the other hand, functional studies show that surface density of Kv4.2 *increases* along this axis (7, 8). Can our modeling and experimental results also account for this relationship?

To address this, we constructed a model of distribution and delivery across a linear multi-compartment dendrite discretized in space. This model is depicted in Figure 6B, where 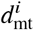 and 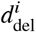 represent microtubule and surface compartments, respectively. 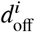 is the rate of microtubule offloading for each 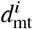, and *f*_*i*_ and *b*_*i*_ denote the forward and backward transport rates along the length of the dendrite. Using our dendritic data (507 mobile puncta in 478 dendrites) to estimate rates, we can infer compatible spatial profiles of surface expression given our experimental data.

**Figure 6:**
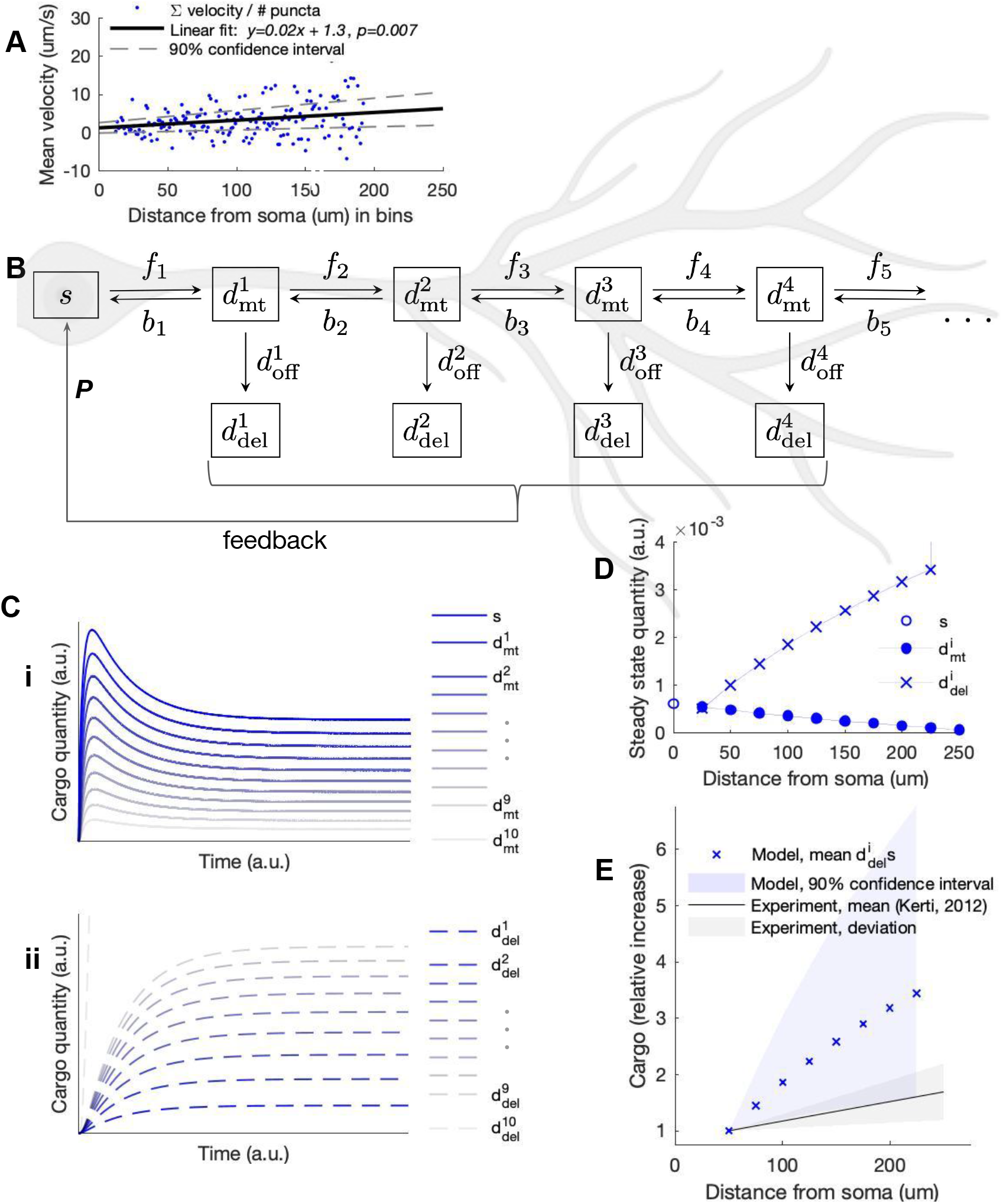
Opposing gradients in static localization and mobile frequency along somatodendritic axis are reconciled with mass-action kinetics. (A): The mean instantaneous velocities for all dendritic puncta are standardized by puncta frequency along the length of the dendrite. A linear tread line is plotted through the data with 90% confidence intervals, indicating a positive (distal) velocity bias that increases with distance from soma. (B): Box diagram of a mass action model of dendritic transport and delivery with feedback. The dendrite is spatially discretized, with each discretization *i* comprising a microtubule 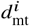 and delivered 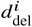 compartment. *f*_*i*_s, *b*_*i*_s, and 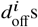 denote rates between compartments. Degradation rates for all compartments are simulated but not depicted. (C): Simulation results depicting cargo quantities in microtubules, 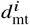 (i), and delivered cargo quantities, 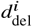 (ii). (D): Steady state concentrations of all compartments. (E): Steady state concentrations of 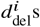 standardized by 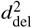 at 50 *μ*m overlaid on equivalently-standardized experimental data of Kv4.2 localization (8).

We considered a dendritic branch extending 250 μm from the soma, which captures typical distances that trafficking and func-tional expression have been characterised. To constrain the steady state concentrations of *mt* compartments 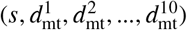, we used experimental values obtained in Figure 5B.

We next computed directional bias in punctal velocity as a function of distance to constrain rates *f*_*i*_ and *b*_*i*_. We averaged the instantaneous velocities of each puncta trajectory in bins by distance from the soma. Mean puncta velocity showed an increasing linear trend with p-value < 0.01, as plotted in Figure 6A with 90% confidence intervals. With a positive y-intercept and slope, the mean punctal velocity is directed distally and increases with distance from the soma. That is, *f*_*i*_ > *b*_*i*_ and *f*_*i*+1_ >> *b*_*i*+1_. The velocities in Figure 6A range from 1.5 to 5.2 μm/s and are scaled according to the spatial discretization of the model to estimate *f*_*i*_s and *b*_*i*_s. The diffusion coefficient 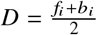 was estimated using Eq. 7, which we assume remained constant throughout the dendritic tree.

As in the previous mass action models, each 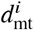 has a corresponding differential equation describing cargo entering and exiting as depicted in Figure 6B. We use steady state analysis to solve for cargo offloading rate 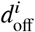 in these equations. We find that a profile of increasing *f*_*i*_s and decreasing *b*_*i*_s with distance from the soma produces an increasing profile of 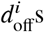. In other words, for cargo with an increasing directional bias such that 0 < *f*_*i*_ − *b*_*i*_ < *f*_*i*+1_ − *b*_*i*+1_ and decreasing *mt* profile, mass action dictates increasing offload rates 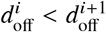 with distance from the soma.

Increasing 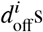 can produce *del* profiles that have the opposite spatial trend to *mt* densities. To demonstrate this, we simulate regulated Kv4.2 production, distribution, and delivery in our model. In the soma, Kv4.2 biosynthesis *P* is regulated by active subunits in *del* compartments, as depicted in Figure 6B. The equation for negative feedback is

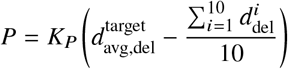

where 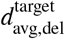 is the target *del* concentration (setpoint), 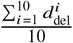 is mean delivered cargo (process variable), and *K*_*P*_ is the sensitivity of the proportional controller. This control loop feedback mechanism is consistent with experimental observations that Kv4.2 expression is regulated as a function of neuron excitability (13–15, 37). Averaging 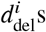 provides a realistic process variable for a global controller in a neuron.

With all rates defined, the result of simulation is depicted in Figure 6C. 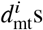 assume a profile similar to that observed experimentally (Figure 5B), with decreasing density with distance from the soma (Figure 6Ci). 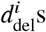 express the opposite profile—increasing density with distance from the soma (Figure 6Cii). Steady state density versus position along the dendrite is plotted in Figure 6D. The increasing 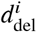 density is notable because localization experiments (8) and, to a larger degree, recordings of A-type current (7) both demonstrate increasing profiles with distance from the soma.

In this analysis, the slope of 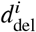 increase (Figure 6D) largely depends on the slope of the mean velocity (Figure 6A) used to constrain the directional bias *f*_*i*_ > *b*_*i*_. In Figure 6E, we plot 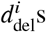 for the linear fit and 90% confidence intervals from our measured directional bias. On the same plot, we shade the reported localization profile of Kv4.2 immunogold-tagged particles from Kerti et al’s 2012 study. Our model constrained by our measured Kv4.2 transport bias predicts an asymmetric profile of delivered Kv4.2 that falls within a standard deviation of localization data. Together these results provide an account of how a previously unexplained and highly organized protein expression pattern can emerge from relatively simple active transport mechanisms.

## DISCUSSION

In this study, we measured mobile Kv4.2-SGFP2 in puncta using recurrent photobleaching while imaging live neurons. Rather than measuring fluorescence recovery as in standard FRAP experiments, we use photobleaching as a tool to remove static fluorescence. This allows for analysis of mobile puncta in bleached regions and basal trafficking of cargo in neurons.

We found substantially higher Kv4.2 subunit trafficking in axons than in dendrites. This surprising result appears to contradict widely documented dendritic expression of this protein. However, just as a satellite photo of car traffic might reveal the highest density of cars on freeways as opposed to parked at a destination, our analysis showed that our measurements are consistent with a mass action model of transport. This implies that increased dendritic demand and surface absorption of Kv4.2 depletes dendritic microtubule-bound subunit density, with axons obeying to opposite trend. Importantly, previous localization studies as well as our own observations reveal a non-negligible density of axonal Kv4.2 (8, 11). With no known presynaptic function, this axonal fraction might be an artifact of mass action, as our study suggests.

Transport kinetics of Kv4.2 puncta in axons qualitatively differed from those in dendrites. Puncta in axons showed increased superdiffusivity, with increased net displacement, increased velocity, and decreased stall time. The opposite is observed in dendrites. This relationship makes sense physiologically, since we assumed that dendrites have a preponderance of sequestration mechanisms that need opportunity to bind cargo and transport it to the membrane. However, increased microtubule offloading is not sufficient to explain the observed differences in kinetics. A random walk with memory better characterizes the experimental distributions for stall fraction and diffusivity. We therefore infer that transport in axons is mechanistically different, with an effective internal state increasing the probability of unidirectional runs.

A number of implicit assumptions are made in our modeling. For instance, microtubule orientation is not considered in mass action or stochastic simulations. Axons have a uniform arrangement of "plus-end-out" microtubules, whereas dendritic orientation is mixed. However, the microtubule motors are also mixed, with both kinesins and dyneins present in all neurites. Our understanding of Kv4.2 interaction with microtubule motors is incomplete, with only Kif17 identified to have a role in subunit trafficking (40). Without a comprehensive understanding of all motors and localization mechanisms, we assume the molecular “tug of war” between motors is equivalent in dendrites and axons, and we implement no additional bias for microtubule orientation. Further investigation can elucidate the observed kinetic differences in axons and dendrites.

We also observed proximal-to-distal trends in dendrites, particularly in puncta frequency and directional bias. When these parameters constrain the rates of a mass action model, the resultant surface concentration of Kv4.2 can account for its well-established, characteristic functional profile (7, 8). A similar increasing profile also exists for hyperpolarization activated cyclic nucleotide-gated (HCN) channels (38). Moreover, a study of HCN channel trafficking and surface expression reveals similar dendritic trafficking dynamics to those reported here but no mention of kinetic trends with distance along the dendrite (39). We suspect that the distance-dependent trafficking parameters observed here are partial contributants to the functional expression profiles of Kv4.2 and HCN channels. Our results reveal that an increasing expression profile does not require an increasing microtubule profile. The increased Kv4.2 expression profile is achievable with a distal directional bias in microtubule transport. By result of mass action, a seemingly complex profile can arise from low magnitude but increasing directional bias. The models used in this study are coarse-grained and not a perfectly detailed depiction of reality. We briefly review the transport and expression mechanisms of Kv4.2 lumped within our models. Kv4.2 interacts with kinesin Kif17, suggesting transport on microtubules. In the absence of Kif17, Kv4.2 fails to localize in dendrites (40). Deletion of a portion of the C-terminus or fusion with myosin Va restricts expression of Kv4.2 to the somatodendritic region (31, 41, 42). Further, KChIPs have been established as auxiliary subunits that promote Kv4.2 exit from the endoplasmic reticulum for surface expression (13, 43, 44). An auxiliary subunit, DPP6, attached to Kv4.2 by a transmembrane domain (45), assists in trafficking Kv4.2 out of the endoplasmic reticulum to the plasma membrane (46). Duménieu et al (47) summarize these results with the following working hypothesis: Kv4.2 is trafficked short distances such as to proximal dendrites or within spines on actin filaments via myosin Va, while long range transport is mediated along microtubules via KChIPs and Kif17.

There are unavoidable methodological tradeoffs between attempting to quantify protein at physiologically low expression levels, and inducing high expression that enables live imaging. We assumed that the transport behavior of the transfected construct Kv4.2-SGFP2 is similar to that of endogenously-expressed subunits. Our results are therefore subject to this caveat. It is possible that transfection of a recombinant construct alters intracellular expression profiles. For this reason, we validated expression profiles by labeling and quantifying both endogenous and transfected Kv4.2 subunits, while using a construct that has been thoroughly compared to endogenous channel (17). We anticipate that our approach can spur future work that will mitigate experimental challenges by designing enhanced fluorescent probes that might be suited to live superresolution imaging. Such methods will be crucial for peering deeper into the logic of intracellular protein regulation.

## AUTHOR CONTRIBUTIONS

A.A.B and T.O. designed the study. A.A.B., L.L., R.S.P., and Y.X.W carried out all experiments. A.A.B. performed numerical simulations. J.G.M. contributed the construct and experiment support. A.A.B., J.G.M., D.A.H., and T.O. designed experiments. A.A.B., J.G.M., L.L., R.S.P., T.O. analyzed results. A.A.B. and T.O. wrote the manuscript. All authors discussed the results and commented on versions of the manuscript.

## ACKNOWLEDGMENTS

We thank Alexander M. Berezhkovskii (Mathematical and Statistical Computing Laboratory, Office of Intramural Research, Center for Information Technology, NIH) for his help with modeling of the modified random walk. We thank Dr. Vincent Schram and Lynne Holtzclaw for their assistance with microscopy performed at the NICHD Microscopy & Imaging Core. We also thank members of the Hoffman lab and the Univ of Cambridge control group for advice and suggestions on this work.

Adriano Bellotti is supported by the National Institutes of Health, the Eunice Kennedy Shriver National Institute of Child Health and Human Development (NICHD), the NIH Oxford-Cambridge Scholars Program, the Gates Cambridge Scholarship Programme, and the UNC School of Medicine MD-PhD Program. Ronald S. Petralia and Ya-Xian Wang were supported by the Intramural Research Program of NIH/National Institute on Deafness and Other Communication Disorders (NIDCD) (Advanced Imaging Core-ZIC DC000081). Timothy O’Leary is supported by ERC grant StG 716643 FLEXNEURO.

